# ISA API: An open platform for interoperable life science experimental metadata

**DOI:** 10.1101/2020.11.13.382119

**Authors:** David Johnson, Keeva Cochrane, Robert P. Davey, Anthony Etuk, Alejandra Gonzalez-Beltran, Kenneth Haug, Massimiliano Izzo, Martin Larralde, Thomas N. Lawson, Alice Minotto, Pablo Moreno, Venkata Chandrasekhar Nainala, Claire O’Donovan, Luca Pireddu, Pierrick Roger, Felix Shaw, Christoph Steinbeck, Ralf J. M. Weber, Susanna-Assunta Sansone, Philippe Rocca-Serra

## Abstract

**Background:** The Investigation/Study/Assay (ISA) Metadata Framework is an established and widely used set of open-source community specifications and software tools for enabling discovery, exchange and publication of metadata from experiments in the life sciences. The original ISA software suite provided a set of user-facing Java tools for creating and manipulating the information structured in ISA-Tab – a now widely used tabular format. To make the ISA framework more accessible to machines and enable programmatic manipulation of experiment metadata, a JSON serialization ISA-JSON was developed.

**Results:** In this work, we present the ISA API, a Python library for the creation, editing, parsing, and validating of ISA-Tab and ISA-JSON formats by using a common data model engineered as Python object classes. We describe the ISA API feature set, early adopters and its growing user community.

**Conclusions:** The ISA API provides users with rich programmatic metadata handling functionality to support automation, a common interface and an interoperable medium between the two ISA formats, as well as with other life science data formats required for depositing data in public databases.

## Findings

### Background

There are many data models and frameworks for describing entities and artefacts of scientific research. The life sciences have pioneered the development and application of ontologies, data standards, and minimum standards for reporting research results. More than ever, the importance of making scientific data FAIR (Findable, Accessible, Interoperable and Reusable) is at the forefront of discourse in the research community [1]. The ISA Metadata Framework, or simply ISA – so named after its constituent key concepts: ‘Investigation’ (the project context), ‘Study’ (a unit of research) and ‘Assay’ (analytical measurement), was born out of initial efforts to create an ‘exchange network test bed’ for a diverse set of digital resources in ‘omics studies. These included public and proprietary databases, as well as free and open source software tools and commercially developed systems. ISA specifies a data model and standard serialization formats for describing experiments in the life sciences.

At the heart of the ISA is a general-purpose data model; an extensible, hierarchically structured model that enables the representation of studies employing one or a combination of technologies, focusing on the description of their experimental metadata (i.e. sample characteristics, technology and measurement types, and sample-to-data relationships). ISA was originally conceptualized and specified as the ISA-Tab format [2,3] for describing life science experiments. Refinements to the underlying data model were later captured in a new JSON-based serialization format, ISA-JSON [4,5] that is specified using JSON schemas [6]. A data model was then abstracted from both the tabular and JSON formats in order to formalize the core concepts and their relationships to one another. The ISA data model is formed around three core concepts: *Investigations, Studies* and *Assays* (Figure 1).

**Figure 1.**
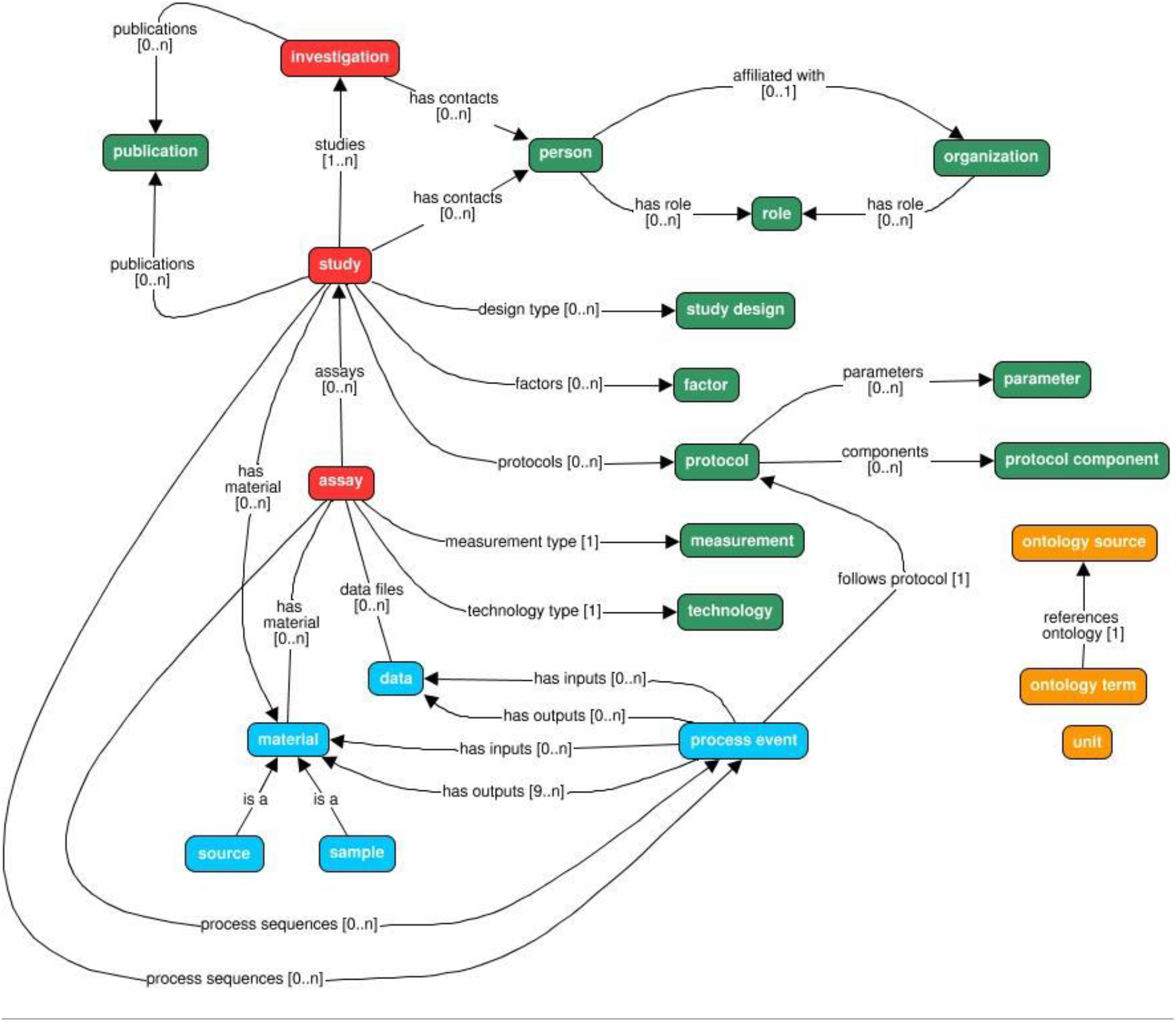
An overview of the ISA data model showing its key constituent classes and their relationships with one another. The model is structured around the concepts of investigation, study and assay (in red). Other model elements exist to qualify these core elements (in green), for example relating investigations and sub-studies with relevant contact persons or related publications; or information about the study design used and any experimental protocols implemented. Experimental workflows are modelled as sequences of process events with inputs and outputs that correspond to biological materials and data objects (in blue). Values can be made explicit by using term annotations or unit declarations that reference published ontologies (in orange).

*Assays* refer to specific tests performed on sample material taken from a subject, or performed on a whole subject, that produces some kind of qualitative or quantitative measurements, which correspond to response variables. Assay metadata describes sample-to-data relationships, grouped by analytical measurement types (e.g. metabolite profiling, DNA profiling) and technologies (e.g. Mass Spectrometry or Nuclear Magnetic Resonance, if the measurement type is metabolite profiling).

*Study* objects hold metadata about the subject(s) under study, including properties about the individual subject(s), any treatments (biological or statistical) applied, and the provenance of sample material back to the original source. Experimental factors relating to the subjects and samples are stored here, allowing the definition of specific study groups in relation to the study independent variables.

*Investigations* contain all the information needed to understand the overall goals and means used in an experiment and are used as high-level objects to group multiple related Study objects. For each Investigation, there may be one or more Studies associated with it; for each Study there may be one or more Assays.

The ISA data model allows us to find scientific experiments of interest by having high-level metadata about the experiments such as what the studies are about and more concretely with descriptors of the Study Design type (a classifier for the study based on the overall study design, e.g. crossover design or parallel group design) and study factors used, e.g. independent variables manipulated by the experimentalist in the study. Furthermore, experiments can be searched by Assay types, on the type of measurements being carried out in the study and the technologies used to perform the measurements.

A key feature in the ISA data model is the use of ontology terms for certain fields in the metadata, supported with the Ontology Annotation and Ontology Source classes. Where appropriate, data model objects can be qualified with ontology terms (Ontology Annotations) that are linked to a declared description of the source of the terms (Ontology Sources). Supporting software tools can therefore populate such annotations from established terminology services such as the NCBI BioPortal [7,8] or the EMBL-EBI Ontology Lookup Service [9,10]. By providing support for discretely identified terms, we enable the ability to search across ISA objects stored in different experiments.

The data model enables experimentalists to describe metadata relating to the provenance of samples and data. This provenance is modelled in the form of directed acyclic graphs (i.e. vertices connected together by edges, but without any loops/cyclic dependencies), which describe the workflow undertaken from subject to sample to the production of data, including associated data acquisition files, analysis data matrices and discoveries being preserved in reusable form.

Full details about the model, as well as specifications for ISA-Tab and ISA-JSON are available from [11].

The supporting open-source ISA tools form a software suite designed to manage the experimental metadata using the aforementioned ISA formats, and more specifically to: (i) customise, describe and validate, following community reporting standards (with the ISAconfigurator, ISAcreator and ISAvalidator components [12] or OntoMaton [13]), (ii) store and browse, locally or remotely (BioInvestigation Index [14] and the ISAexplorer [15]); (iii) submit to public repositories (ISAconverter [12]); (iv) analyse with existing tools (Risa) [16]; and (v) publish data alongside the article. The adoption of these Java-based and web-based software tools has helped grow the ISA community of users, characterised by those listed in the ISA Commons [17].

As the original ISA software suite does not provide a programmatic interface, we have developed the ISA API to position the ISA framework as an interoperable and open platform [18]. We also recognized a trend of growing enthusiasm by end users for writing software code. This trend is perhaps most evident in statistical programming platforms such as R, Python and MatLab, and also greatly helped with the uptake of interactive programming environments such as Project Jupyter [19].

The aim of the ISA API is to reproduce much of the user-facing functionality afforded by the ISA software suite and to additionally enable interoperation between software systems that produce and consume ISA formats and other machine-readable data formats, as illustrated in Figure 2. The ISA API is written in Python, which has a high uptake by non-programmers and a rich open source community ecosystem of supporting software packages, including many statistical and bioinformatics ones. The software code is available on GitHub [20] under the Common Public Attribution License Version 1.0 (CPAL-1.0) and the software package is available via the Python Package Index (PyPI) [21] and Bioconda [22] under the moniker: `isatools` [23,24].

**Figure 2.**
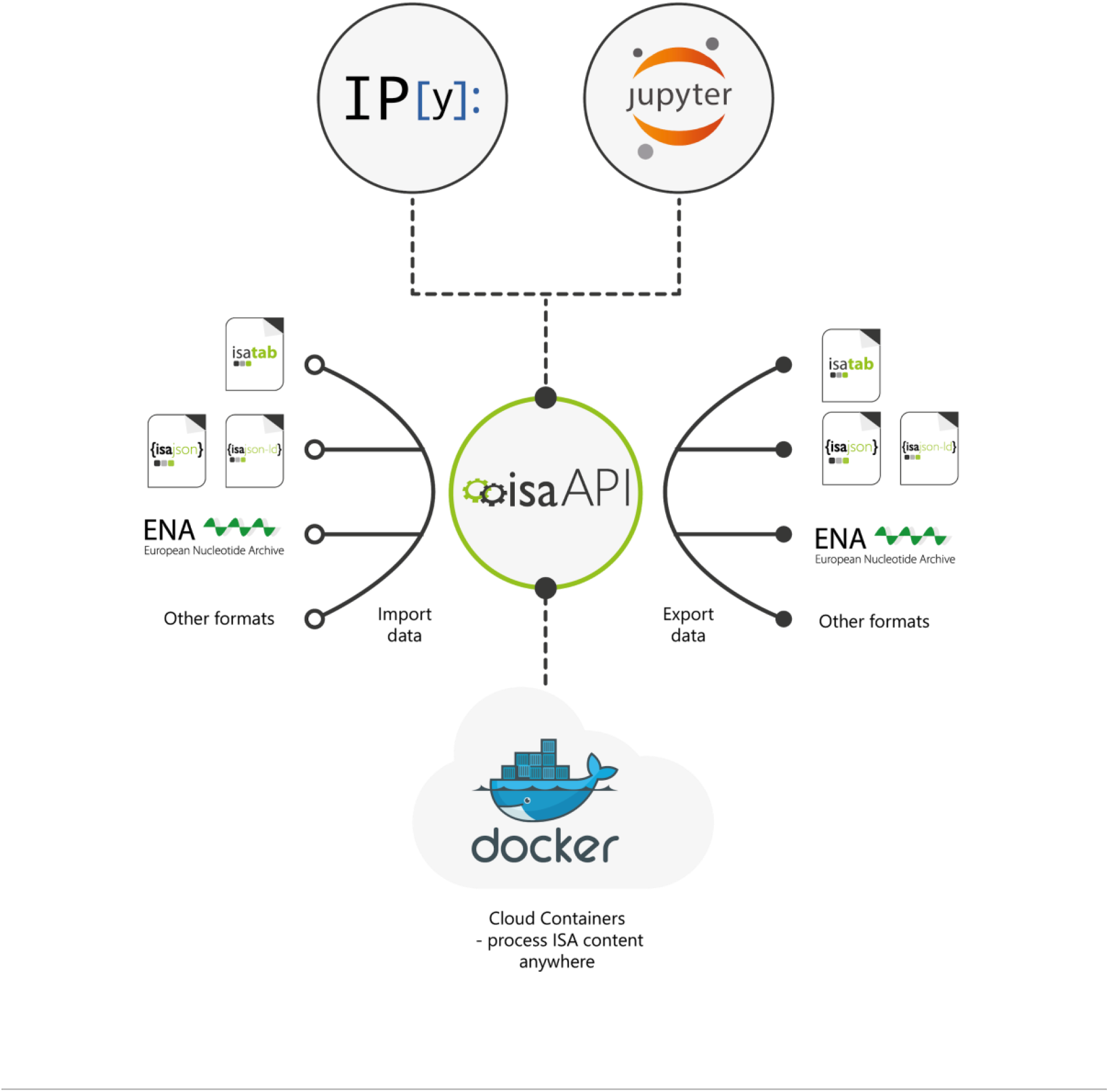
ISA API and its interactions with other software ecosystems and data formats. Apart from running in the Python interpreter, the ISA API can also be accessed through the iPython, Jupyter and Docker. It also supports interoperation with other systems through standardised machine-readable data formats.

### Related work

Developed in early 2011, the `Bio-Parser-ISATab` project [25] is, to the best of our knowledge, the first effort to create an ISA-Tab parser outside of the ISA tools ecosystem for programmatic manipulation of ISA metadata. While `Bio-Parser-ISATab` is written in the Perl programming language, the first Python project to implement an ISA-Tab parser with scripts found in the `biopy-isatab` project [26] was developed as part of an early version of `bcbio` [27] later in 2011. Both efforts focused only on reading ISA-Tab.

Recent efforts to create ISA programming libraries have focused on reading and writing of ISA-Tab files: AltamISA [28] was developed in Python with the aim for strict validation and error handling when handling ISA-Tab files; and a Java library, isa4j [29], has been developed for high-performance writing of ISA-Tab metadata.

ISA API provides a reference implementation of ISA-Tab support and additionally supports other formats and features as described in the following section.

## Implementation

The ISA API is written in the Python 3 programming language, and therefore part of the wider Python ecosystem of software tooling which is very popular in the bioinformatics and data science communities [30]. The ISA API can immediately be imported and used alongside other Python packages in custom Python programs, or it can be used interactively via iPython’s interactive shell [31] and in Jupyter Notebook web applications. It can be used in cloud infrastructure that supports Docker through the container image `isatools/isatools` published in DockerHub. The ISA API can be also exposed through RESTful APIs as demonstrated by the MetaboLights Labs Web service interface and the MetaboLights Online Editor Web service [32].

## ISA API features

The ISA API provides a range of features that are summarized in this section.

### Support for all ISA formats

The ISA API provides the solution to simplify access and generation of ISA metadata now that the ISA framework supports more than one on-disk data format – i.e., ISA-Tab and ISA-JSON. By using the ISA API, software can support full read, write and validation transparently on all current and future ISA formats without any additional complexity. The ISA-Tab and ISA-JSON document validators included in the ISA API provide reference implementations by which to also validate documents generated by other systems that claim ISA compatibility [17].

### Support for other bioinformatics formats

The ISA API also supports related domain formats used by bioinformatics communities and which are usually technology-centric (e.g. mass spectrometry, sequencing, DNA microarrays). This feature is a consequence of the ISA framework’s support for multi-modality datasets (e.g. multi-omics datasets). The ISA API supports the import and export of MAGE-TAB [33,34] and SampleTab [35,36], import of metadata from mzML [37,38] and nmrML [39,40], and exports to SRA-XML [41,42] and Workflow4metabolomics [43].

### Export metadata for submission to public repositories

The ability to export metadata to a range of formats means that software using the ISA API can prepare metadata submissions to public data repositories. The ISA API supports export of ISA metadata to submission-ready formats for public repositories such as the EMBL-EBI ArrayExpress [44,45], European Nucleotide Archive [46,47]s, and MetaboLights [48,49]. Support for submission to additional databases will be developed in future.

### Multiple modes of handling ISA metadata

The ISA API is designed to support multiple modes of reading and writing ISA. Metadata reading and writing can be done natively with ISA-Tab folders, ISArchive zip files, and ISA-JSON files or strings. ISA metadata can be manipulated after loading using an object-oriented Python class model. Direct conversion between ISA formats is also supported.

### Extensible and coherent object-oriented class model

Metadata objects are represented in the ISA API using an object-oriented class model, providing a format-agnostic in-memory representation. This in-memory model is based on the ISA Abstract Model, which was extracted from the original ISA-Tab specification to formalize the concepts used in the ISA framework. It describes the steps in an experimental workflow as a directed acyclic graph (DAG), which in the ISA API is expressed naturally as connected Python class objects. For example, metadata relating to the assay description is held in an `Assay` object that is in-turn contained by a `Study` object and its relevant attributes, which is finally contained by an `Investigation` object – thus reflecting the hierarchical nature of the ISA framework concepts. All ISA concepts are modelled as classes at a granular level giving software developers full control to extend the API’s capabilities and build new features. Details of the class model used in the ISA API are published in the ISA Model and Serialization Specifications 1.0 [4].

This functionally rich object-based representation completely avoids many of the complexities associated with handling ISA metadata as tables – e.g., joining data from the sample and assay tables, handling multiple table rows per edge of the DAG. While in memory, the experimental metadata can be programmatically processed and manipulated. Moreover, the ISA API’s object-based representation is a powerful decoupling tool. First, it decouples the client application from the specific ISA file format being handled. Second, it decouples the input and output metadata formats. When metadata is read, the format-specific parser creates the standardized in-memory representation, which can be edited and manipulated; when writing, a format-specific serialization routine traverses the objects to extract the data structure according to the target ISA format. Writing the ISA API object model to ISA-Tab requires converting experimental workflow graphs to tabular formats. Such work is described in the `graph2tab` library by Brandizi et al [50]. However, in that work, the authors only describe cases of unidirectional transformations from graphs to tables; ISA API implements bidirectional transformations between each of the two ISA formats and ISA content expressed as Python objects that includes a representation of DAGs. ISA content can also be authored by directly creating Python objects, a brief example of which is shown in Figures 3 and 4.

**Figure 3.**
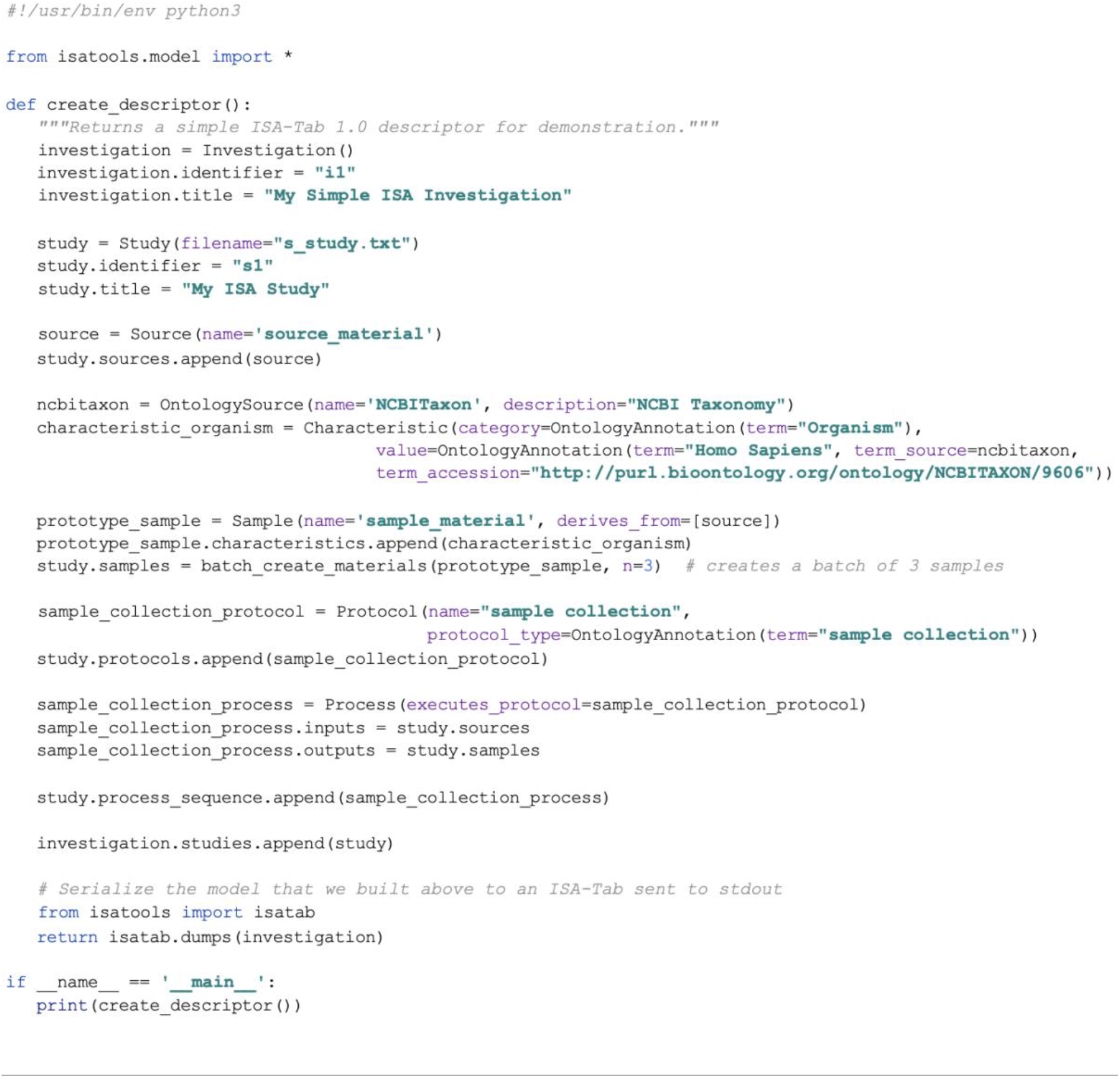
Example of a Python script using the ISA API’s class model objects to programmatically construct metadata about a study and serialize it to ISA-Tab. This script first creates the Investigation and Study structures that store general metadata about the experiment being described. Next, a source material object is created, and three sample materials. These are connected as inputs and outputs respectively to a sample collection process, forming a workflow from source to samples. This is then added to the study’s ‘process sequence’: a container for experimental process event descriptions. Finally, the composition of the model objects is serialized as ISA-Tab to the standard output. Scripts such as these can form part of a larger Python software program, or executed directly from the command-line, to automate the construction of ISA metadata descriptors.

**Figure 4.**
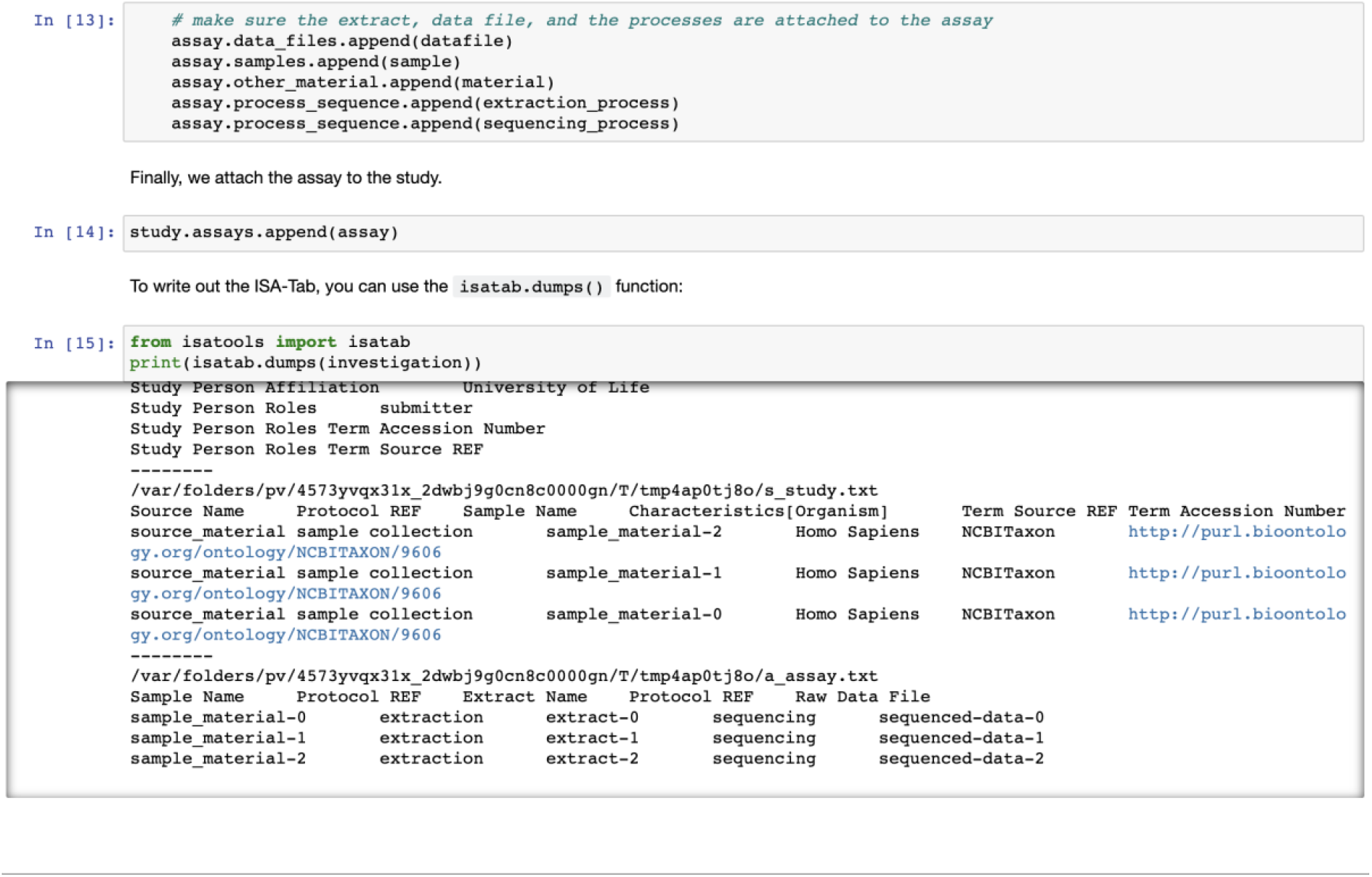
Example Jupyter Notebook using the ISA API to use ISA class objects to programmatically construct metadata about a study, using similar code to that shown in Figure 3. Being Python-based, ISA API can integrate with any notebook environment that supports Python kernels including Jupyter Notebook, JupyterLab and JupyterHub, and proprietary notebook environments such as Google Colab [51], Microsoft Azure Notebooks [52], and Amazon’s SageMaker [53]. A set of Jupyter notebooks detailing how to use ISA-API key functionalities is available on GitHub [54].

### Querying over ISA content

Common use cases for ISA metadata involve gathering samples and data files produced by assays that satisfy certain selection criteria. The criteria are objects based on sample characteristics, processing parameters, and study factor combinations (treatment groups). When ISA metadata is loaded as in-memory model objects, this representation can be queried using native Python code, for example by filtering using list comprehensions with conditional logic. The ISA API also provides helper functions, in the `isatools.utils` package, to query and fix possible encoding problems in ISA content. For example, in ISA-Tab files, a common issue is when ISA content is encoded in such a way that some branches in experimental graphs converge unexpectedly which represents the intended workflow incorrectly. The ISA API provides specific functions to detect such anomalies (in the `detect_isatab_process_pooling` function) and to fix them (in the `insert_distinct_parameter` function). Details of how to query over ISA content can be found in the ISA API online documentation [55].

### Semantic markup

An important feature that is embedded into the class model is the support for ontology annotation objects to better qualify metadata elements, contributing to making them Findable, Accessible, Interoperable, and Reusable (FAIR) [56]. The ISA API implements the `OntologyAnnotation` class that is used extensively throughout. To express quantitative values, units are implemented with the `Unit` class, which itself can be qualified with an `OntologyAnnotation` object. Populating annotations is aided by a network package in the ISA API for connecting to and querying the Ontology Lookup Service (OLS) [9]. JSON-LD context files complementing the ISA-JSON schemas mapping into schema.org [57] or OBO Foundry based resources [58] are also available.

### Assisted curation of study metadata

Studies published using ISA formats are increasingly commonplace, many of which are produced in ISA-Tab using the ISAcreator (for example, NASA GeneLab [59,60] and MetaboLights [48]), and many others produced either by-hand (using a spreadsheet or text editor) or output by third-party systems (for example, COPO [61], FAIRDOM/SEEK [62,63] and SCDE [64,65]). There are sometimes cases of invalid ISA-formatted content appearing in public databases or reported via bug reports to the ISA development team. To help with this, the ISA API includes features to assist with metadata curation of ISA-formatted studies. Beyond using the validators contained in the API, additional features include checking for possible structural problems in the DAGs that describe the provenance of samples and data files (for example, where converging edges are detected but are not expected) and correcting them; re-situating incorrectly placed attributes (e.g. recasting incorrectly labelled `Factors` as `Parameters` or `Characteristics`); and batch fixing such issues. This work has resulted in the creation of curation functions, which have been applied to the content of the MetaboLights database as a means to improve the quality and consistency of ISA archives. These validators also form the basis of the current MetaboLights extensive online validations coupled to their submission infrastructure to validate the metadata for missing values, consistency and sufficiency.

### Assisted creation of study metadata

The ISA API *create* module contains a set of functions and methods exploiting study design information to bootstrap the creation of ISA content [66]. Initially created to support factorial designs, this component is currently being extended to provide support for longitudinal studies and repeated treatment designs. This work in progress will be refined in future versions of the ISA API.

### Documentation

Extensive documentation is available as a Jupyter Book that includes examples presenting the various capabilities afforded by the ISA API [55]. The Jupyter Book includes information about how to install and use the `isatools` Python package, details about the ISA data model, and source code examples of using ISA API as Python scripts and Jupyter notebooks. For more information about the features listed here, interested readers are invited to view the ISA tools API documentation for detailed and up-to-date information.

## Early adopters

The ISA API has a number of early adopters that demonstrate its value as an open platform to the ISA framework. The following section details how these groups rely on our python library in their data management tasks.

The MetaboLights metabolomics database [48] is using the ISA API to load, edit, and save ISA-Tab as part of its next-generation data curation interface. MetaboLights has redeveloped its user and data curator interface building on the latest Web technologies and using the ISA API to convert from ISA-Tab to ISA-JSON for further processing. The developers take a service-oriented approach exposing various features for editing and updating ISA content through a RESTful API [32].

The PhenoMeNal (Phenome and Metabolome aNalysis) e-infrastructure for molecular phenotype data analysis has developed a set of containerised microservices (using Docker) that utilise ISA API’s converters, validators, and study creation features [67]. The PhenoMenal Virtual Research Environment (VRE) is based on the Galaxy workflow platform [45] through its integration with Kubernetes [68]. Through PhenoMeNal, a suite of tools has been developed to integrate the ISA API with Galaxy [69]. The tools have been published as a Galaxy workflow since the Dalcotidine release of PhenoMeNal to demonstrate a study metadata preparation and pre-submission pipeline to the MetaboLights database (see Figure 5).

**Figure 5.**
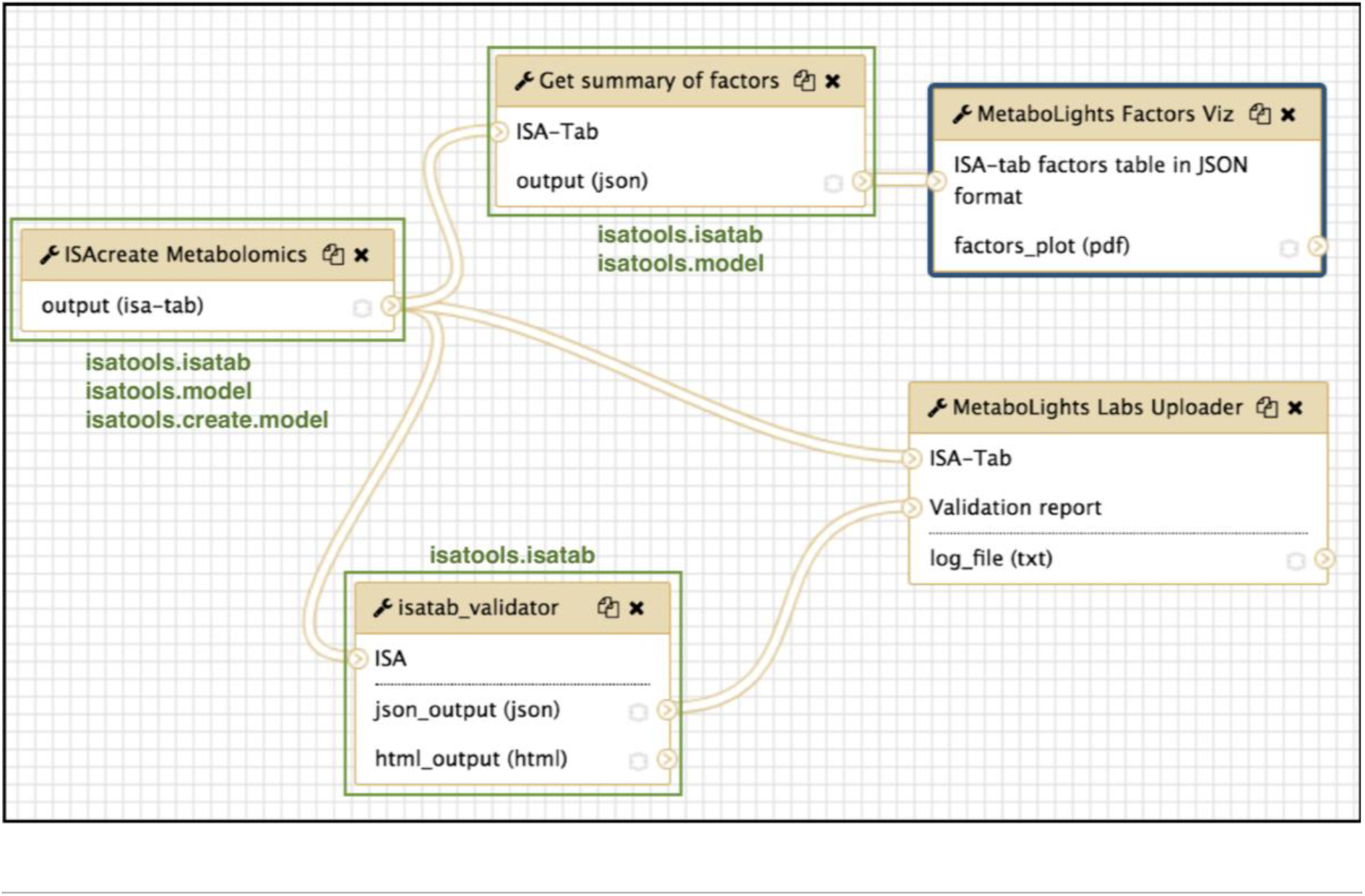
The ISA Create-Validate-Upload workflow, published as part of the PhenoMeNal platform Dalcotidine release. The workflow takes a user-configured study plan and creates an ISA-Tab template ready for the experimentalist to use in their study. The ISA-Tab then goes through two paths of processing: (1) a summary of study factors according to the study design is extracted and then visualized as a parallel sets plot; and (2) the ISA-Tab is validated, and if valid a pre-submission request is made by uploading the ISA-Tab to MetaboLights Labs. A preparatory study accession ID is then issued by MetaboLights if accepted. Parts of the workflow that directly use the ISA API are highlighted in green along with the packages used.

The Galaxy workflow platform includes native support for ISA formats by using the ISA API in its datatype implementations of ISA-Tab and ISA-JSON, paving the way for Galaxy tools to consume and produce ISA content as first-class Galaxy datasets.

The Collaborative Open Omics (COPO) platform [70] supports ISA-JSON as one of its metadata formats and uses ISA API’s SRA exporter to allow COPO users to publish studies directly to the European Nucleotide Archive. COPO configures its back end and front end for brokering various data types, including omics data, through consuming ISA configurations. The ISA configurations inform how COPO should present the data preparation wizard to the user, and how it stores it in its database in a variation of ISA-JSON. COPO stores study metadata as JSON where it conforms to the ISA-JSON schemas and adds extra metadata relating to its operation within the COPO database. By supporting ISA-JSON, COPO lends itself to easily leveraging the ISA API to export data to formats supported by the API. Still within the plant science community, a conversion from BrAPI [71] to ISA has been developed along with MIAPPE compliant ISA configurations [72,73]. The functionality will be integrated along with other community conversion components in future releases of the ISA API.

## A stable and growing user community

Since its first release, ISA API has grown its user base steadily with active contributions from the community as an open source project as evidenced by bug reports, feature requests, and code contributions. While it is difficult to measure the uptake of free software, ISA API has been available since 2016 via PyPI, which collects download statistics for all packages it hosts [74]. There are documented drawbacks to using these PyPI download statistics, such as historical data issues and systemic irregularities such as caching of downloads that can cause undercounting, so we view these statistics as indicative of how the ISA API project has progressed rather than as an accurate measure. These statistics also do not include installations directly from the source code repository on GitHub or from Bioconda.

After its initial release year, we observed a maintained and slight year-on-year growth from 2017 to 2019 of around 27,000 annual downloads via PyPI (Figure 6), which we believe demonstrates that the ISA API has built up a stable and active user base.

**Figure 6.**
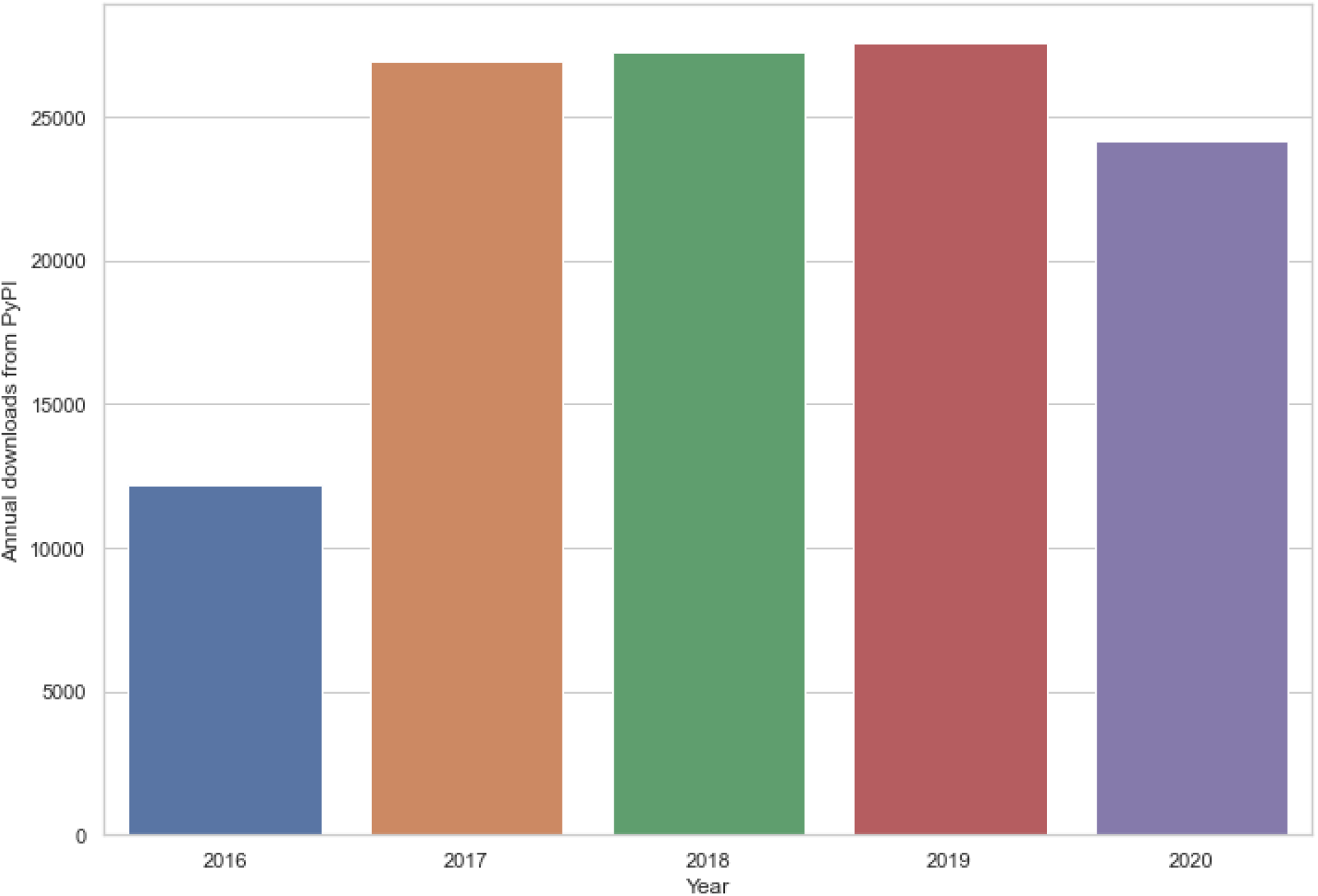
Download statistics the `isatools` Python package from PyPI from 21 February 2016 (first release of ISA API, v0.1) up to 24 October 2020 (after release v0.11). Note that the total number of downloads for 2020 was incomplete at the time the data was collated. Sub-figure (6a) shows a bar chart depicting the annual total downloads of `isatools`. Sub-figure (6b) shows a line chart depicting the monthly total number of downloads of `isatools` with major releases of the ISA API indicated with vertical dashed lines.

**Figure 7.**
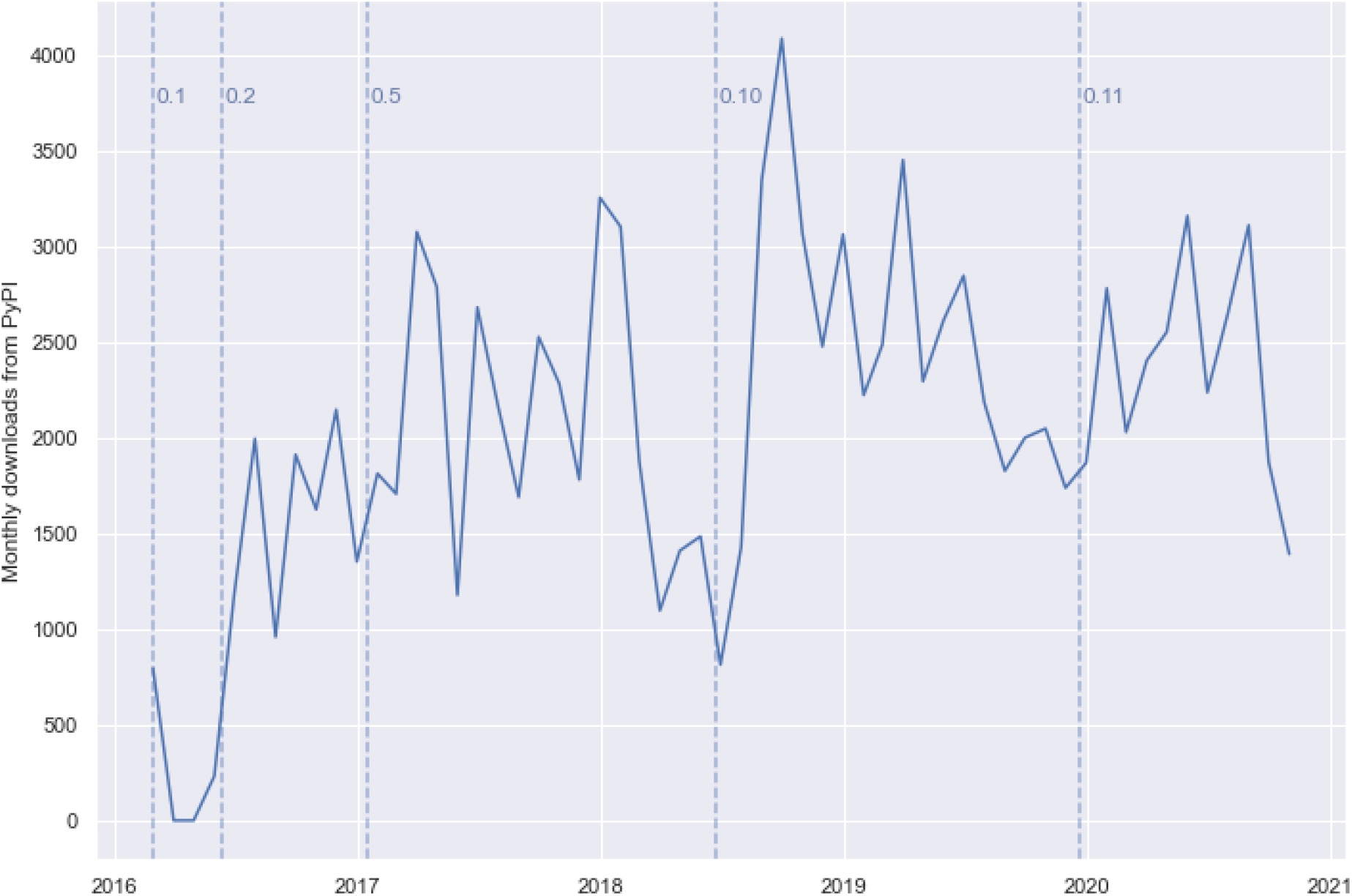

When looking at a more granular picture of the download statistics for the `isatools` package, on a month-by-month basis, we observe spikes in downloads on PyPI after each major release. We believe this indicates that the ISA API user community continues to be active and is using the platform in production systems by migrating to its newest updates soon after they are made available.

## Conclusions

The development of the ISA API represents a major step forward in making the ISA framework open and interoperable, enabling better handling of experimental metadata, and supporting a diverse user community. By providing a programming interface, rather than a graphical user interface, the ISA API provides a platform for simplified and consistent integration of the ISA framework in new and existing software, promoting the production of structured metadata by agents and the development of data management solutions that can be FAIR by design. The availability of an ISA implementation that is easily used by other software also reduces the likelihood of independent implementations that may not strictly adhere to the ISA format specifications. The object-oriented class model provides a format-agnostic interface for client software allowing software that uses the ISA API to automatically support all ISA file formats, while also providing a common basis on which to build data format parsers and serializers more easily. Should external implementations of the ISA formats be required, by providing reference implementations of ISA-Tab and ISA-JSON parsers, serializers and validators, software developers have the infrastructure in which to extend the API or build interoperation of ISA with other software systems. As Python code, it can also easily integrate into existing bioinformatics and data science platforms such as Galaxy and Jupyter. Importantly, the object-oriented DAG model presented by the ISA API offers a more intuitive way to reason about and process study metadata than as ISA-Tab tables, simplifying the work of developers and enabling the development of richer metadata processing applications. The ISA API is available as free and open-source software which will improve the dissemination and application of the ISA metadata framework.

## Methods

### Community-led requirements

The ISA metadata framework and its supporting tools were developed to support requirements of the life science research community and its requirements are elicited in collaboration with a wide range of stakeholders, such as those represented in the ISA Commons [17]. This approach continues with the development of the ISA API where, through direct research funding, the project was initially driven by requirements of metabolomics research (via the H2020 PhenoMeNal project [67]) and the plant sciences community (via the UK BBSRC COPO project [70]). Development has also been guided by collaboration workshops (for example [75] and [76]). As an open source project from its outset in 2015, the ISA API openly encourages grass-roots contributions via its GitHub project, where we receive continuous feedback, feature requests, and source code contributions from third parties.

### Model-driven engineering

The ISA API was developed using a model-driven engineering (MDE) approach [77] that initially focused heavily on formalising the concepts that define the ISA metadata framework. This required close analysis of the only specification of ISA concepts available documented in the ISA-Tab 1.0 specification [2] and the de facto reference implementation of ISA-Tab in the Java source code of the ISA software suite [12], as well as critical discussions within the ISA Working Group. One of the early goals of the ISA API project was to implement support for a JSON serialization format, ISA-JSON [4]. A model specification was drafted as a set of JSON schemas, the JSON community’s response to implementing features reminiscent of XML schemas. This was then formalised as the first data agnostic specification of ISA. From the ISA-JSON schemas, we then used the `warlock` Python package [78] to draft an object-oriented (OO) Python class model (via Python’s dynamic object creation facility) that was highly synchronised with the JSON schemas. This OO class model was then implemented concretely (to provide object reflection/introspection that dynamic objects lack) to use as the primary representation of ISA metadata in Python. Data format parsers and serializers were then developed to load data into these Python objects at run-time. The ISA API implements support interoperation between formats such as ISA-Tab, ISA-JSON, SampleTab, MAGE-TAB and SRA-XML by using these ISA Python objects.

It is important to note that the ISA data model is not a model of ISA-Tab; rather, ISA-Tab is an implementation of the data model. The representation of ISA concepts in a tabular form is specific to the serialization format, where ISA is not about tables, rows and headers. ISA metadata is about the encapsulated semantics about the life science experiments being described. However, in developing the class model, ISA-Tab parser and serializer it was realised that some ISA-Tab concepts were required in the class model to support serialization.

For example, the ISA-Tab table file names are not strictly part of the ISA data model but we need to be able to specify these to support writing ISA-Tab files. When writing to ISA-JSON these file name attributes are ignored as the JSON implementation is more closely aligned to the model. Therefore, the ISA API’s implementation of the model should be considered as only very closely aligned with the model specification rather than a perfect mapping.

### Test-driven development

When developing new features and fixing bugs, the ISA API development team predominantly follows a test-driven development (TDD) approach [79]. This means that when new features are requested an automated unit test is implemented first to capture the feature specification, then work is done to develop the new feature to satisfy the test. Similarly, when a bug is reported via the project’s GitHub issue tracker, an automated regression test is implemented first to capture the problem specification before developing the bugfix. Automated unit tests are implemented using the `unittest` package and run with the `nose` and `tox` testing frameworks. Source code contributions to the GitHub project are automatically tested on multiple Python versions using the continuous integration service Travis CI. ISA API’s most recent release at time of writing (v0.12) uses 631 automated unit tests.

### Download statistics

The download statistics for the `isatools` package hosted on PyPI shown in Figure 6 are collected by the Linehaul statistics daemon [80] that logs downloads of all Python packages on PyPI and pushes the data to a public dataset [74] on Google BigQuery [81]. The all-time statistics about the `isatools` package were retrieved from this public dataset on 24 October 2020 at 14:20 UTC+2 with the following SQL query via the BigQuery interface:

~~~
SELECT TIMESTAMP_TRUNC(timestamp, WEEK) week
, COUNT(*) downloads
FROM ‘the-psf.pypi.downloads*’
WHERE file.project=‘isatools’
GROUP BY week, file.version
ORDER BY week
~~~

The result of this query was downloaded as a CSV file containing a timestamp of the week-ending each weekly period of downloads, the `isatools` version number, and the total number of downloads for each period and version. This data was then aggregated by month and by year before being visualized using the Seaborn statistical data visualization package [82]. Data about the version release dates of ISA API were collected from the ISA API GitHub repository [20].

## Availability of supporting data

The all-time weekly download statistics data for the `isatools` package on PyPI up to 24 October 2020 14:20 UTC+2 and the code used to plot the charts used in Figures 6a and 6b in this manuscript are available in a Code Ocean capsule [83]. The data and code is available under CC BY 4.0 and MIT License respectively.

## Availability of source code and requirements

- Project name: ISA API
- Project home page: http://github.com/ISA-tools/isa-api/
- Operating system(s): Platform independent
- Programming language: Python
- Other requirements: Python 3.6+
- License: CPAL-1.0
- biotools: isa_api
- RRID: SCR_020980

## Declarations

## List of abbreviations

API: Application Programming Interface
BBSRC: Biotechnology and Biological Sciences Research Council
BrAPI: Breeding API
COPO: Collaborative Open Plant Omics
DAG: Directed Acyclic Graph
EMBL: European Molecular Biology Laboratory
EMBL-EBI: European Bioinformatics Institute
GUI: Graphical User Interface
H2020: Horizon 2020
ISA: Investigation, Study, Assay
ISA-JSON: ISA JavaScript Object Notation format
ISA-Tab: ISA Tabular format
JSON: JavaScript Object Notation
JSON-LD: JavaScript Object Notation for Linked Data
MAGE-TAB: MicroArray Gene Expression-Tabular format
MIAPPE: Minimum Information About a Plant Phenotyping Experiment
mzML: Mass spectrometry Markup Language
NASA: National Aeronautics and Space Administration
NERC: Natural Environment Research Council
NCBI: National Center for Biotechnology Information
nmrML: Nuclear magnetic resonance Markup Language
MDE: Model-driven engineering
OLS: Ontology Lookup Service
OO: Object-oriented
PhenoMeNal: Phenome and Metabolome aNalysis
PyPI: Python Package Index
REST: Representational state transfer
RDF: Resource Description Framework
SCDE: Stem Cell Discovery Engine
SQL: Structured Query Language
SRA-XML: Sequence Read Archive-eXtensible Markup Language
TDD: Test-driven development
VRE: Virtual Research Environment

## Consent for publication

Not applicable.

## Author’s Contributions

Conceptualization: P.R-S, S-A.S.; software-lead: D.J.; software-supporting: M.I., P.R-S, A.G-B, K.C., A.E., K.H., M.L., T.N.L., A.M., P.M., V.C.N., L.P., P.R., F.S., R.J.M.W.; investigation-usage: D.J.; visualization-usage: D.J.; writing-original draft: D.J.; writing-review and editing: D.J., P.R-S., K.C., R.P.D., A.E., A.G-B, K.H., M.I., M.L., T.N.L., P.M., V.C.N., C.O., L.P., P.R., F.S., C.S., R.J.M.W., S-A.S.; funding acquisition: R.P.D., C.O., S-A.S, C.S.

## Acknowledgements

We thank members of the ISA Working Group, ISA Commons, H2020 PhenoMeNal project, UK BBSRC COPO project, attendees of the 2016 “Hack-the-Spec - ISA as a FAIR research object” workshop hosted by the Oxford e-Research Centre, UK, attendees of the 2016 “China-UK Data Dissemination in Metabolomics (CUDDEL)” workshop hosted by GigaScience, Hong Kong, and the EMBL-EBI MetaboLights team for guidance and feedback on the development of the ISA API.

## Author information

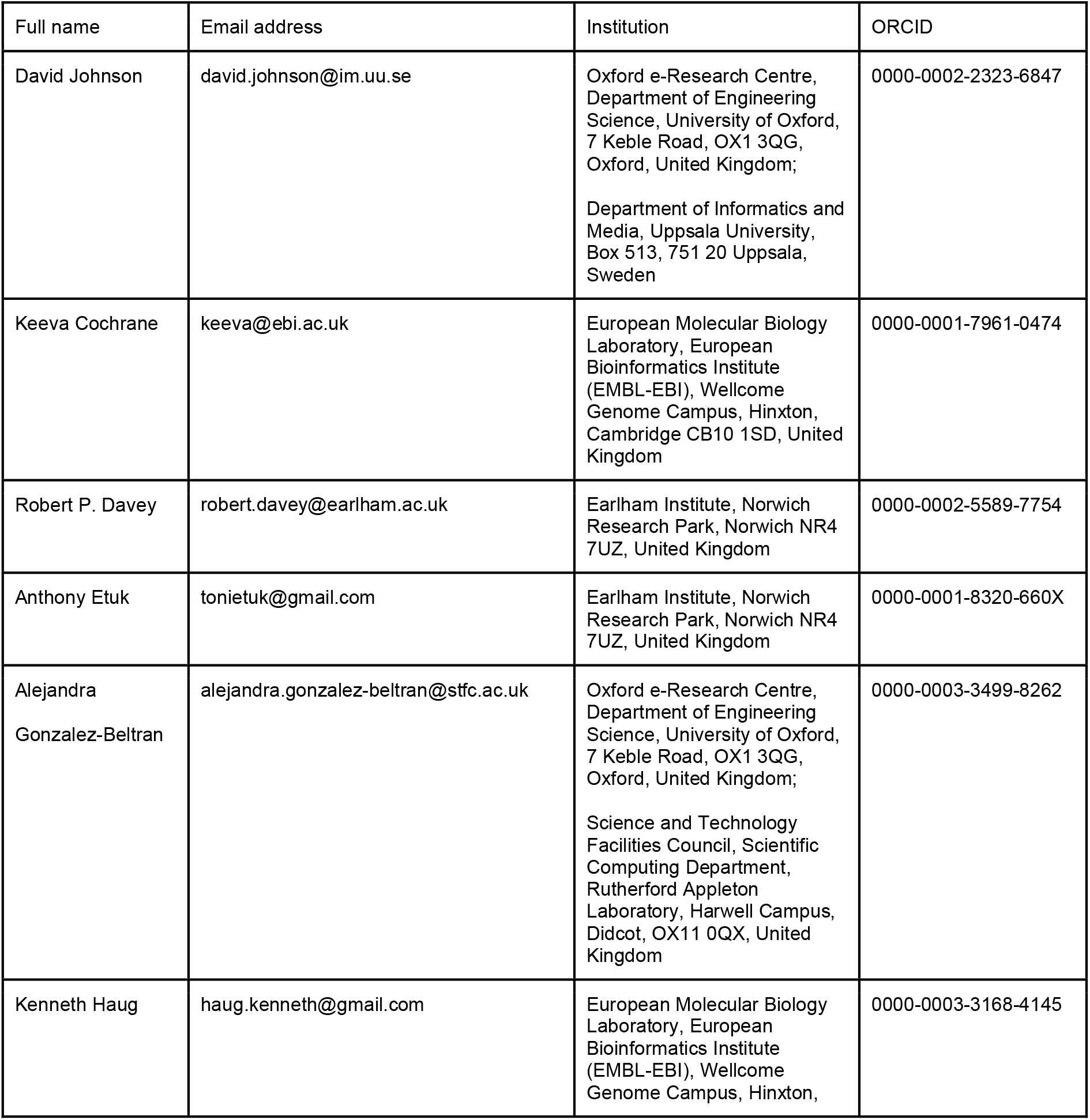
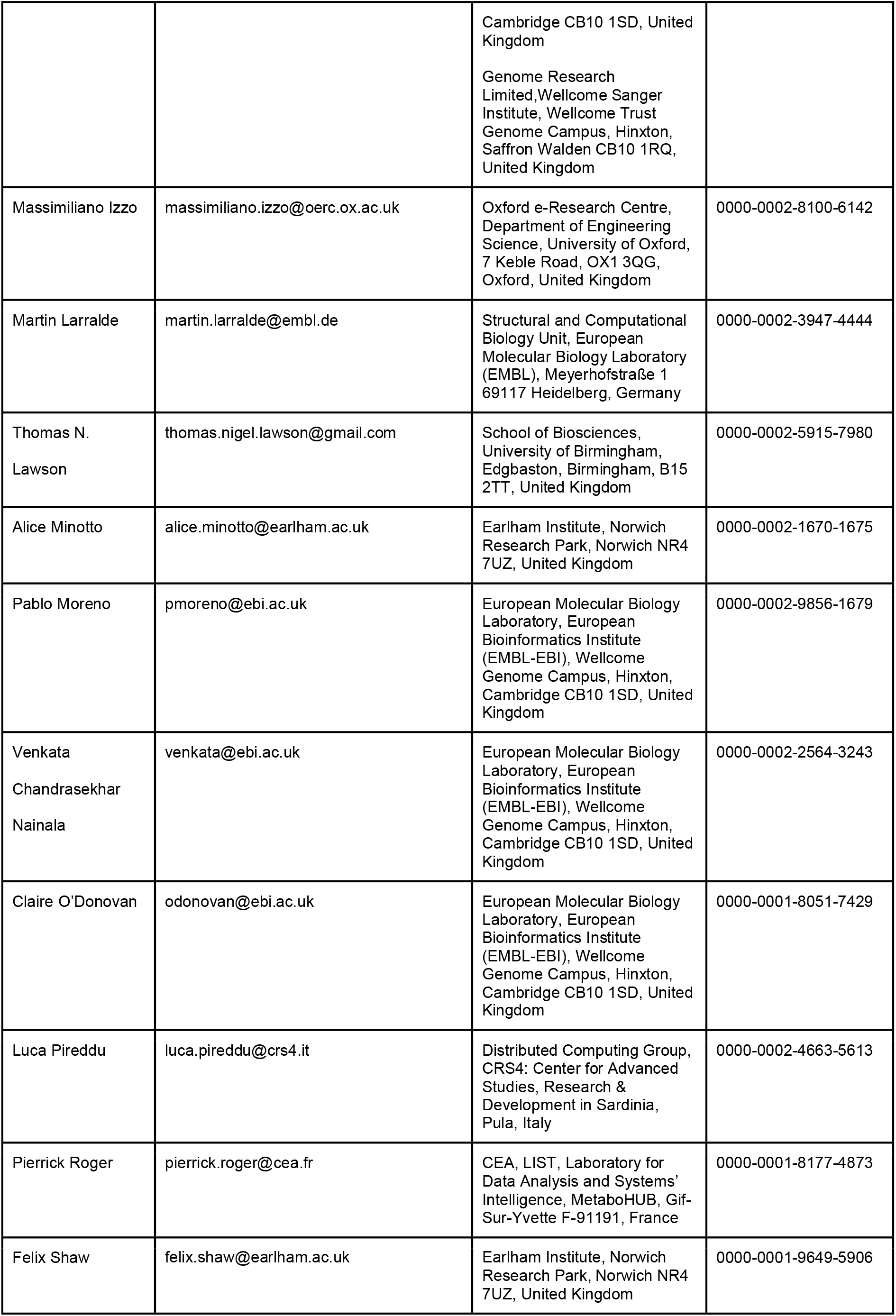
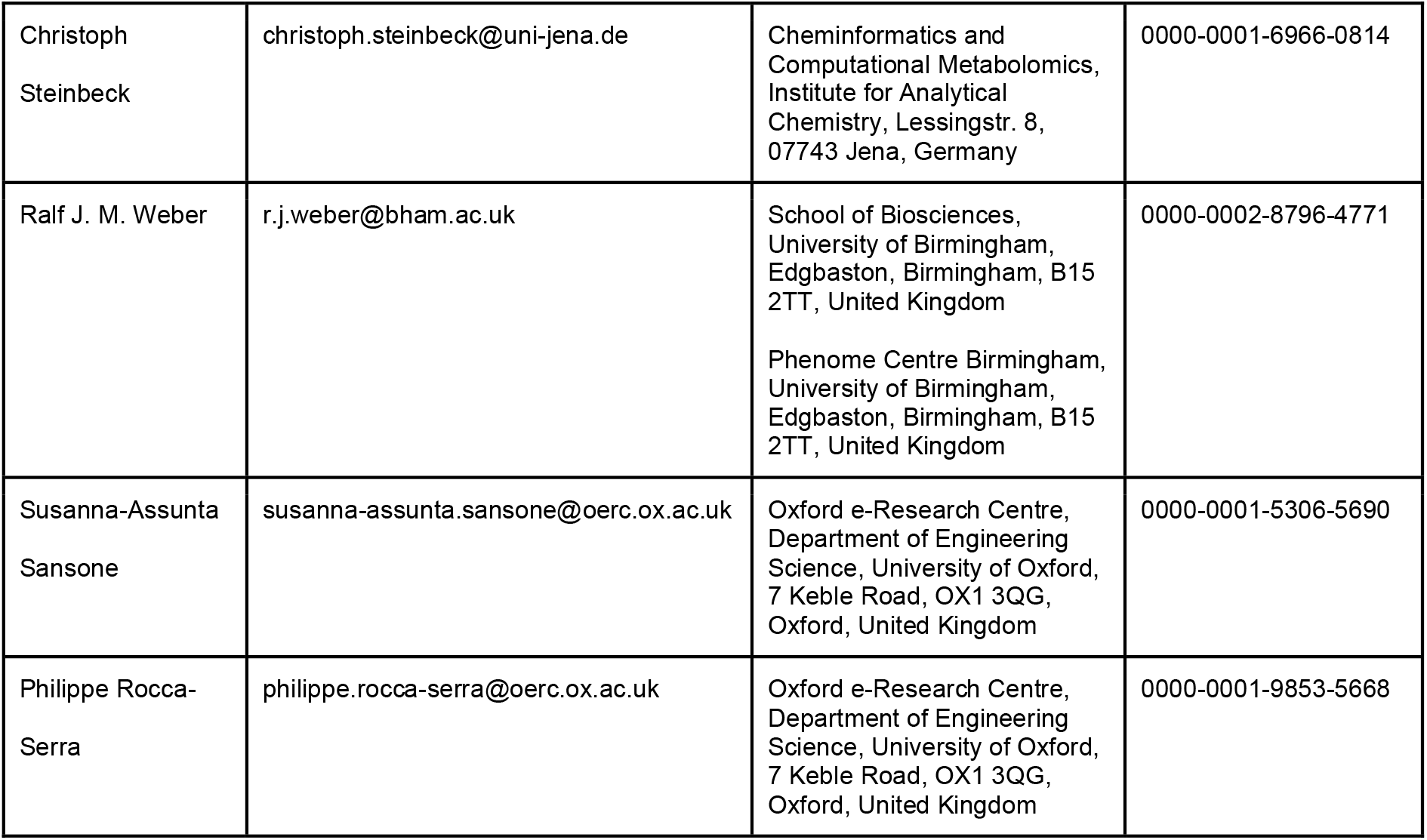

